# Large-scale integrated single-cell analysis reveals molecular subtypes and microenvironmental remodeling in cutaneous T cell lymphoma

**DOI:** 10.64898/2026.10.22.761138

**Authors:** Ryan J Reis, Eric Mou, Vincent Liu, Patrick M Brunner, Nicholas Borcherding, Ali Jabbari

**Affiliations:** Medical Scientist Training Program, University of Iowa, Iowa City, IA, USA; Department of Dermatology, University of Iowa, Iowa City, IA, USA; Cancer Biology Graduate Program, University of Iowa, Iowa City, IA, USA; Division of Hematology, Oncology, and Blood and Marrow Transplantation, Department of Internal Medicine, University of Iowa, Iowa City, IA, USA; Holden Comprehensive Cancer Center, University of Iowa, Iowa City, IA, USA; Department of Pathology, University of Iowa, Iowa City, IA, USA; Department of Dermatology, Icahn School of Medicine at Mount Sinai, New York, NY, USA; Department of Pathology and Immunology, Washington University, St. Louis, MO, USA

## Abstract

Cutaneous T cell lymphoma (CTCL) comprises a clinically and molecularly heterogeneous group of skin-homing non-Hodgkin lymphomas for which robust biomarkers remain limited. Prior single-cell RNA sequencing (scRNA-seq) studies have provided important insights into CTCL biology, but many of these prior studies have been limited by small sample sizes and focused on niche questions of interest. Additionally, many studies have suffered from pseudoreplication, resulting in inflated test statistics and reduced external validity. Here, we assembled and integrated scRNA-seq data spanning 244 unique samples and 132 patients across CTCL, healthy control (HC), and benign inflammatory dermatoses (BID) conditions, generating the most diverse atlas of CTCL to date. Following identification of malignant cell populations and heterogeneous, within-sample gene expression alterations, we applied pseudobulk aggregation and robust linear mixed-effects modeling to nominate recurrent changes across this highly heterogeneous patient cohort. Interrogation of the skin microenvironment identified significant compositional and cell type-specific expression changes in CTCL lesions as compared to control lesions and across clinical stages of disease. Finally, we used unbiased clustering to identify stable, molecularly defined subtypes of disease that were predictive of outcome in an independent dataset. Together, this study provides a rigorous analysis of the most diverse scRNA-seq dataset of CTCL ever compiled, nominating specific alterations in cell type composition and gene expression across malignant cell and microenvironmental populations with significant clinical implications.

## Introduction

CTCL encompasses more than a dozen unique types and subtypes of non-Hodgkin, skin-tropic lymphomas and lymphoproliferative diseases [1]. Collectively, these subtypes are defined according to unique histopathological, molecular, and clinical features. Remarkably, despite longstanding efforts to understand the genetic basis of these diseases, few recurrent single-nucleotide variants (SNVs) have been identified [2, 3]. Rather, copy-number variations (CNVs) and other structural and complex genomic alterations appear to be more common [3–8]. However, these events are heterogeneous and their effect on downstream cellular behavior is incompletely understood. Collectively, these limitations have culminated in an incomplete understanding of CTCL pathogenesis and progression and continued unmet clinical needs of these patients.

Single-cell RNA sequencing (scRNA-seq) has emerged as a powerful technology for studying disease processes at a molecular and cellular resolution [9–11]. As such, numerous studies have applied scRNA-seq technologies to study CTCL. These studies have identified unique and heterogeneous expression changes across malignant cells and their environments. However, many of these studies have been limited by sample size and focused on specific patient populations of interest, limiting interpretation of these results in a broader context. Additionally, many groups have reported results using cell-level statistical testing, which treats each cell as an independent data point. This form of pseudoreplication can introduce bias into the reported results, inflate test statistics, and fail to replicate in external cohorts [12, 13].

To address these challenges, we assembled and integrated the most diverse dataset of CTCL scRNA-seq data to date. Using sample-specific cell-level testing, we first revealed the highly heterogeneous nature of CTCL across patients. We then implemented a statistically rigorous approach pairing sample-aware pseudobulk aggregation with linear mixed-effects modeling to account for patient- and batch-specific variation [14–16]. Using this approach, we identified robust expression changes in CTCL cells as compared to their phenotype-matched counterparts, robust molecularly defined subtypes of CTCL within the skin compartment, and changes within the tumor microenvironment (TME) as they relate to disease state, clinical stage, and the identified subtypes. Finally, we validated the prognostic significance of our findings in an external cohort of patients [17].

## Results

### An integrated multi-cohort atlas of CTCL

In total, 244 sequencing results from 132 patients and 18 experimental cohorts were integrated into a single dataset (Fig 1a) [18–34]. Following quality control and filtering, approximately 1.57 million cells were retained for downstream analysis. Uniform manifold approximation and projection (UMAP) visualization of the final integrated dataset colored by cell type (Fig 1b) as well as sample, cohort, diagnosis, and degree of T cell receptor (TCR) clonal expansion binned by proportion of each sample, referred here as clonesize, (Fig 1c) are presented. TCR sequencing was completed for 208 of the 244 total samples and largely corroborated the assigned cell types, with TCR assignment 97.9% specific to the annotated T and innate lymphoid cell (T/ILC) population. Differential gene expression analysis was performed across the identified cell types. The top three genes specific to each annotated cell type were selected using a combined rank score of the average log2 fold change (log2FC) and percent difference in cell expression (delta expression) and are presented (Fig 1d). To better reflect the original biology of the samples, those sequencing results originating from the same biological tissue and artificially separated prior to library prep were merged, herein termed a sample. Finally, per-cell type proportions were calculated for each sample and confirmed high fidelity of the annotated populations across the included tissue sites (Fig 1e).

**Figure 1:**
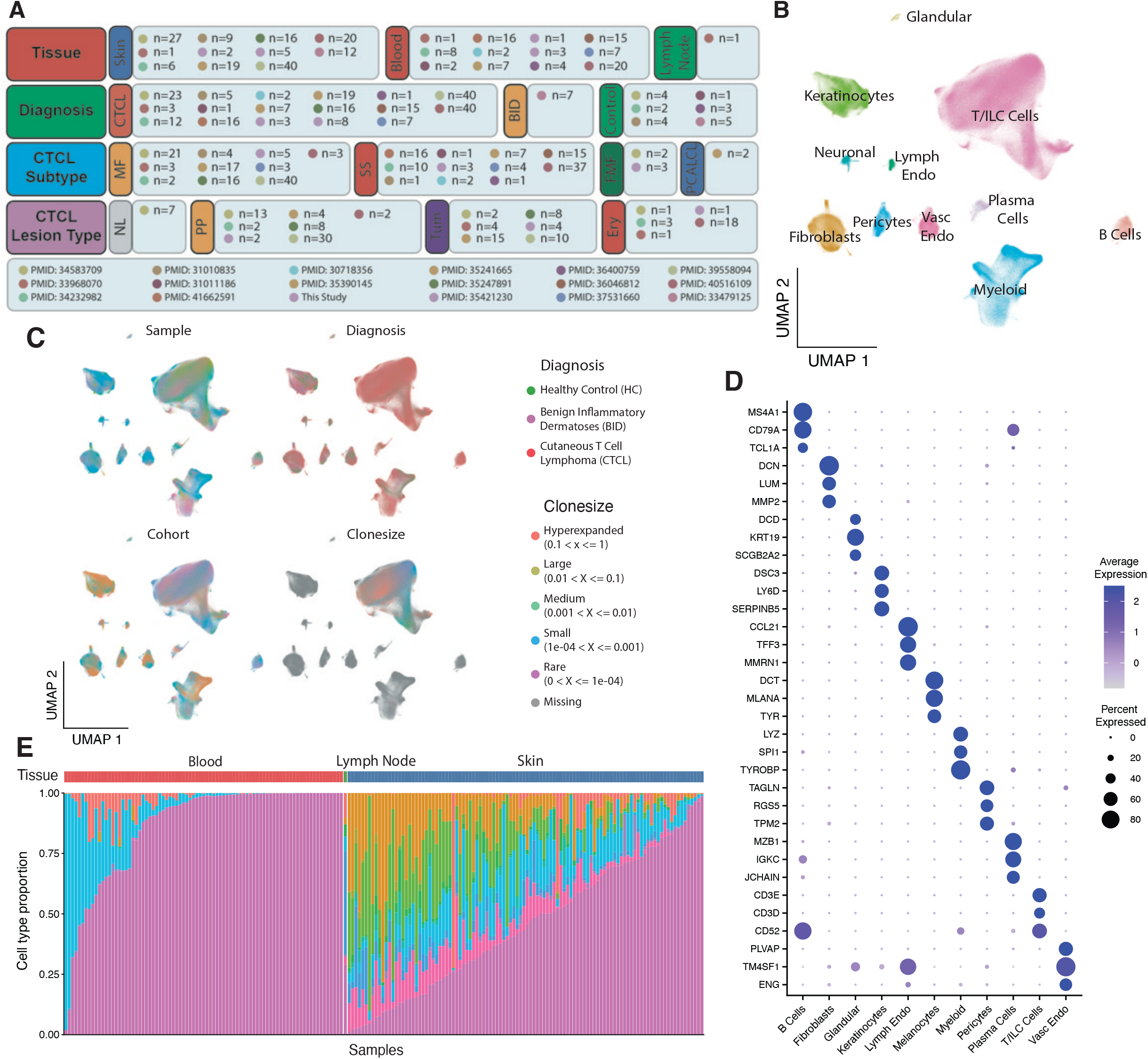
A cross-tissue, integrated atlas of CTCL, healthy, and benign control disease states. A. An overview of the scope and distribution of samples across tissue sites, major diagnoses, CTCL subtypes, and CTCL lesion types across all data sources. B. UMAP representation of the final filtered and integrated object, colored by annotated cell type. C. UMAPs of the final object colored by sample, cohort, diagnosis, and degree of T cell clonal expansion, as determined via TCR sequencing. D. A dot plot of the top marker genes across each of the annotated cell types. E. A stacked bar chart representing the relative proportion of each annotated cell type within each sample, split by tissue site. T/ILC, T and innate lymphoid cells; Lymph Endo, Lymphatic Endothelial; Vasc Endo, Vascular Endothelial

### Malignant cell identification requires a multi-modal, curated approach

Following integration and identification of major cell populations, we sought to specifically distinguish malignant CTCL cells from background, normal T/ILC populations. Upon isolating the identified T/ILC population from other cell types in the dataset, we assessed the distribution of the clonesize (Fig 2a) and expression of common T/ILC phenotypic markers (Fig 2b). While detection of expression of *CD4* was limited, a known issue with scRNA-seq technology, we noted significant clonal expansion across regions consistent with the expected dominant immunophenotype of these patients, most commonly alpha-beta (αβ) TCR+, CD3+, CD4+, CD8- [1, 35–37]. However, due to the heterogeneity of our samples and sparsity of single-cell data, we determined that we were unable to reliably discern CTCL cells from the benign populations using the integrated expression results alone.

**Figure 2:**
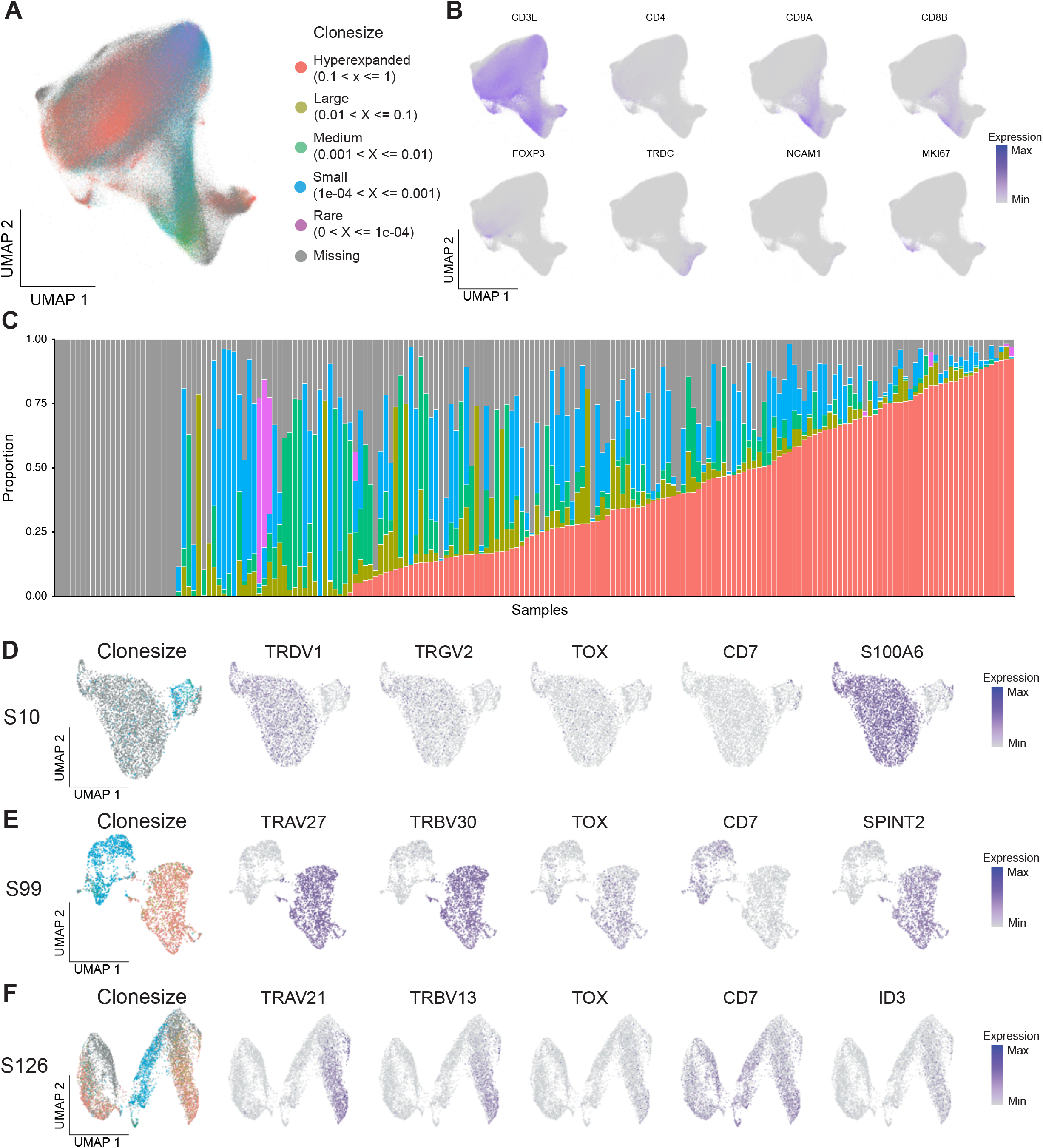
Clonotypic and transcriptomic characterization of the T/ILC compartment. A. UMAP of the subset T/ILC compartment, colored by degree of clonal expansion. B. Feature plots of canonical phenotypic T cell and ILC gene markers. C. Stacked bar charts representing the proportion of each assigned clonesize per sample. D. UMAP visualization of S10, colored by clonesize and expression of top TCR, CTCL, and sample-specific marker genes. E. UMAP visualization of S99, colored by clonesize and expression of top TCR, CTCL, and sample-specific marker genes. F. UMAP visualization of S126, colored by clonesize and expression of top TCR, CTCL, and sample-specific marker genes. S10, Sample 10; S99, Sample 99; S126, Sample 126

Although contested, CTCL is largely believed to be a monoclonal expansion of mature T cells [24, 38, 39]. Because of this, many researchers have relied on paired TCR sequences to assess the clonality of their cells using scRNA-seq technology. However, we found that relying on these TCR sequences alone was insufficient to reliably discern these populations with high specificity, due to the lack of paired data across all samples and the sparsity of recovered data, limitations commonly encountered in scRNA-seq technologies (Fig 2c). Beyond these issues, we found several cases where TCR-based approaches alone could not reliably separate CTCL cells from their benign counterparts. Specific examples of this include a rare case of gamma delta (γδ)-derived CTCL (S10, Fig 2d), case of dual TCR alpha chain usage, a known phenomenon in normal T cell biology, (S99, Fig 2e), and a sample harboring two hyperexpanded T cell populations, of which one was determined to be a responding CD8+ T cell clone (S126, Fig 2f) [40]. We therefore adopted an integrative approach using both gene expression and TCR-seq data, where available, to assign malignant and benign labels on a per-sample basis following reprocessing of these individual samples. While both data modalities individually struggle to reliably discern these populations at the cellular level, this integrative, cluster-based approach proved successful.

### Cell-level differential gene expression testing recovers canonical markers of disease and patient-level heterogeneity

Once malignancy status was annotated across all cells and samples, the combined CTCL and benign T/ILC object was reprocessed. A UMAP was generated using the unintegrated reduced dimensions and revealed strong intra-sample structure, particularly within the malignant cell fraction (Fig 3a,b), consistent with unique inter-patient expression profiles. In an attempt to first identify top gene expression changes in CTCL across the aggregate cohort of samples, a Wilcoxon rank-sum test was performed, comparing all CTCL cells against all benign T/ILC populations. While this proved to recover many of the commonly described alterations in gene expression, notably the downregulation of *CD7* and *GSTP1* and overexpression of *TOX*, *KIR3DL2*, *PLS3,* and *AIRE* in CTCL cells relative to their benign counterparts (Fig 3c), this approach is unable to recover sample-specific heterogeneity and may mask rare expression changes [18, 41–44].

**Figure 3:**
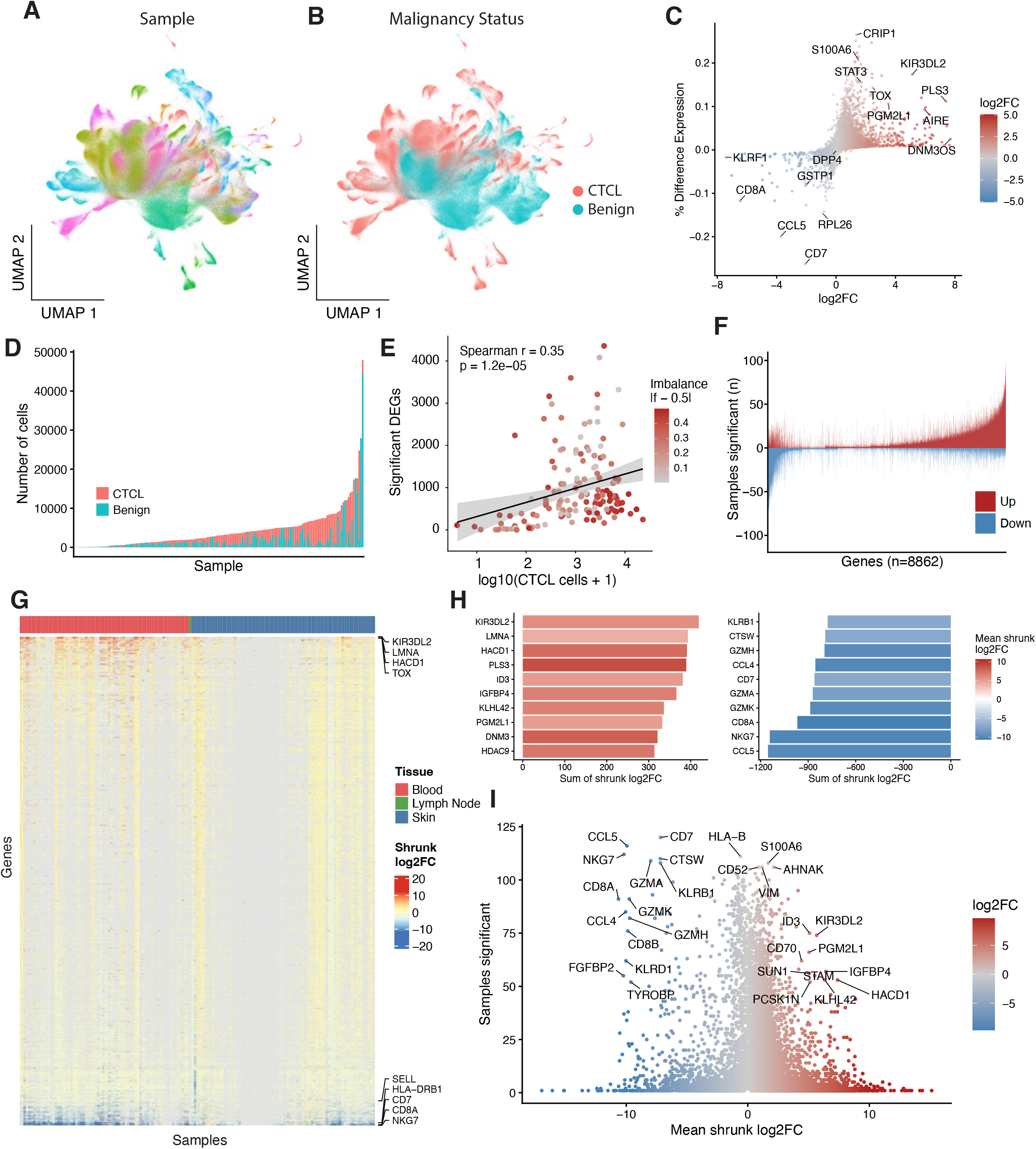
Identification of alterations in gene expression in CTCL. A. UMAP visualization of the unintegrated, reprocessed T/ILC population, colored by sample. B. UMAP visualization of the unintegrated, reprocessed T/ILC population, colored by annotated malignancy status. C. Scatter plot of the pan-sample, cell-level DEG testing comparing CTCL to benign T/ILC cells. D. Bar chart of the number of cells and relative malignancy status of them, within each sample. E. A scatterplot of the results of the sample-specific DEG testing, comparing the number of significant genes detected to the number of cells and colored according to class imbalance of the tested populations. F. Bar chart showing the number of samples for which each gene was found to be significantly increased (positive, red) or decreased (negative, blue). G. A heatmap of the shrunk log2FC values of the recurrent (>10%) significant genes detected per sample, ordered by tissue and total summation across samples. H. Bar charts depicting the top 10 up and down DEGs across all aggregated samples, as calculated and ordered by summation of shrunk log2FC values. I. Volcano plot depicting the summarized results of the aggregated, per-sample DEG testing as a function of the number of samples a gene was found to be significant in and the mean shrunk log2FC value across significant samples. CTCL, Cutaneous T cell lymphoma

We therefore performed differential gene expression analysis across each CTCL and benign fraction within each sample, individually. As expected, both the number of cells and relative proportion of each fraction varied widely across samples (Fig 3d) and were associated with the number of differentially expressed genes (DEGs) detected (Fig 3e). However, this approach proved successful in identifying sample specific alterations, with a total of 148,430 significant gene-sample DEG events identified across the 151 tested samples. Notably, many of these significant results were heterogeneously detected across samples, with the direction of change also varying widely across many events (Fig 3f, Supplementary). To identify top recurrent expression changes, we shrunk and aggregated the log2FC values of those significant genes across all samples, retaining those genes identified in at least 10% of samples (Fig 3g). Summation of shrunk log2FC values across samples recapitulated many of the commonly described genes with increased expression in CTCL, although many of the top downregulated genes likely represent differences due to normal T/ILC phenotypic variation and the identification of a dominant CD4+ CTCL phenotype across this patient cohort (Fig 3h, i). For example, many of the top downregulated genes include *CD8A*, *CD8B*, and many other known activating and cytotoxic genes expressed in CD8+ T cell and ILC populations such as *NKG7*, *CCL5*, *GZMK*, and *CTSW*.

### Pseudobulk mixed modeling identifies recurrent CTCL gene expression changes

To improve nomination of gene expression changes across samples while reducing biases introduced by cell-level testing procedures, we next implemented a pseudobulk approach coupled with linear mixed-effects modeling, in order to properly model unwanted sources of variation across cohorts [12, 14, 16, 45]. Briefly, pseudobulk aggregation combines expression results across multiple cells within a sample, most often using a simple summation, as depicted (Fig 4a). By taking this approach, we both reduce sparsity and variation of expression introduced by single-cell sequencing, while also mitigating pseudoreplication in subsequent testing [12, 14, 16]. Unlike many prior efforts, we also identified and subset non-T regulatory (Treg) CD4+ benign cells, specifically, to use as the comparator population, reducing the effect of normal T cell phenotypic variation on these results.

**Figure 4:**
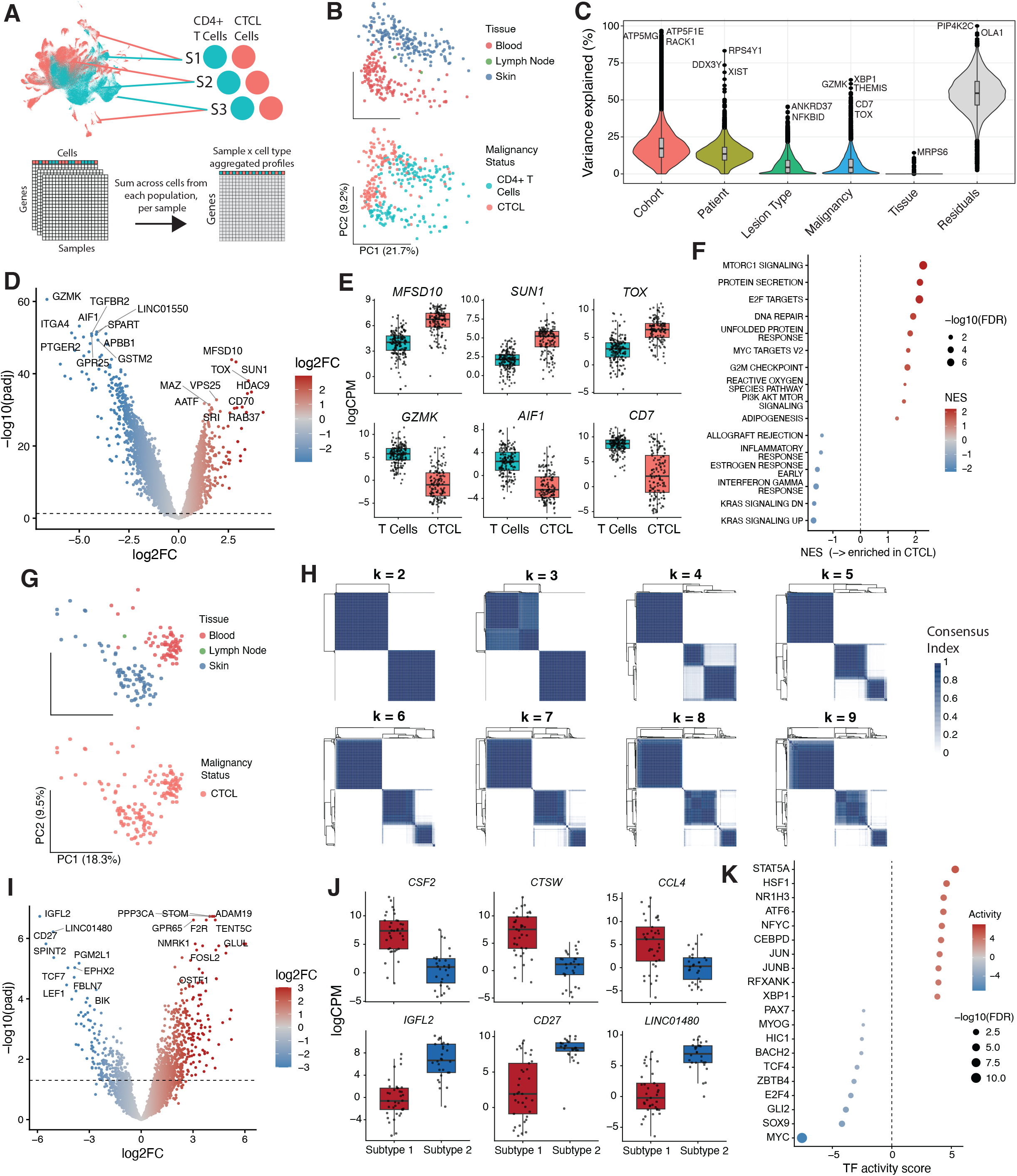
Comparison of gene expression across CTCL and identified molecular subtypes. A. A schematic representing the process of cell type-aware pseudobulk aggregation using the tested populations of interest. B. PCA plots showing the pseudobulk aggregated profiles across the top two principal components, colored by tissue site and malignancy status. C. Violin plots showing the results of variance partitioning of gene expression within these pseudobulk profiles across select metadata of interest. D. Volcano plot of the results of the linear mixed-effects modeling, comparing CD4+ phenotypic CTCL samples to the benign counterparts. E. Box plots of logCPM values of select top markers, split by malignancy status. F. Dot plot of the GSEA results from the malignancy DEG testing. G PCA plots showing the pseudobulk aggregated profiles of CTCL cells across the top two principal components, colored by tissue site and malignancy status. H. Heatmaps depicting the consensus clustering index of each pairwise sample across consensus clustering results. I. Volcano plot of the results of the linear mixed-effects modeling, comparing CD4+ Subtype 1 vs Subtype 2, as determined by consensus clustering assignment. J. Box plots of the logCPM values of select top markers from the subtype DEG testing, split by assigned CTCL subtype. K. Dot plot of the inferred TF scores as determined by TF inference analysis of the subtype DEG results. CTCL, Cutaneous T cell lymphoma; logCPM, log counts per million; NES, Normalized enrichment score; FDR, false discovery rate

Pseudobulk aggregation was completed within the identified CTCL and benign CD4+ T cell populations, and the resultant aggregated matrix was normalized and filtered for downstream analysis. Principal component analysis (PCA) was performed, identifying tissue and malignancy status as major sources of variation in the batch-corrected pseudobulk matrix (Fig 4b) [12]. We next assessed the influence of select metadata of interest on gene expression, nominating top variable genes across each (Fig 4c) [16]. Formal linear mixed-effects modeling was performed, modeling malignancy status and tissue as fixed effects and patient and cohort as random effects. This identified 7,550 significant DEGs between the populations, including numerous lowly-expressed genes that would otherwise be filtered out using standard single-cell approaches (Fig 4d). Top candidate downregulated genes include *GZMK*, *AIF1*, *LINC01550*, *ITGA4*, *APBB1,* and *TGFBR2*, with *CD7* ranking as the 20^th^ most downregulated gene by adjusted p-value (padj). Notably, none of these top genes have been nominated as canonical features of CTCL outside of select contexts, as loss of *CD7* has been [46]. Conversely, top upregulated genes include *SUN1*, *MFSD10*, *TOX*, *HDAC9*, *CD70*, and *VPS25*. While increased TOX expression in CTCL is well described, many of these other top candidate genes have only been implicated in CTCL in limited contexts, if at all [22, 47–50]. Select boxplots of these top genes are presented in Figure 4e. Finally, gene set enrichment analysis (GSEA) of these results against the MSigDB Hallmark (MSigDB: H) gene sets demonstrated significant enrichment of pathways related to mTORC1 and MYC signaling as well as E2F targets, DNA repair, and the unfolded protein response, among others (Fig 4f) [51]. Interestingly, genes associated with classical immune effector and inflammatory pathways were relatively reduced in the CTCL profiles, suggesting that a substantial fraction of inflammatory signals within these lesions may arise from background normal T cell populations.

### Consensus clustering reveals recurrent subtypes of disease within the skin compartment

We next asked whether pseudobulk CTCL profiles could be used to identify recurrent molecular subtypes of disease, independent of known metadata. PCA of the CTCL aggregated matrix revealed strong tissue-driven variation (Fig 4g), but distinct clusters beyond that were not appreciated via PCA nor non-linear transformation. Rather, variable gene identification and consensus clustering successfully identified two robust clusters, annotated as Subtype 1 and Subtype 2, of disease within the skin compartment and across a range of k values (Fig 4h). To assess the robustness of these results, samples were annotated according to their cluster assignment in the k=4 clustering, and linear mixed-effects modeling was performed to compare these two populations. This testing confirmed the results of our unbiased clustering, identifying 859 significant genes between them (Fig 4i). Select top markers from each identified subtype are presented (Fig 4j).

To identify the underlying transcription factors (TFs) and pathways associated with each of these distinct molecular subtypes, we next performed GSEA and TF inference analysis. GSEA revealed increased expression of genes related to inflammatory, cytotoxic, and T cell activation pathways in Subtype 1. Subtype 2 showed increased expression of MYC, oxidative phosphorylation, and cell cycle-related genes (Supplementary). Additionally, TF inference analysis confirmed the unique phenotypes of these samples, with Subtype 1 samples showing significant inferred activity of STAT5A, HSF1, NR1H3, and ATF6, among other inflammatory and cytotoxicity-related TFs; Subtype 2 was found to have significantly increased inferred activity of MYC, SOX9, BACH2, and other central-memory and non-canonical TFs (Fig 4k). While these results are largely consistent with the tissue-resident memory/cytotoxic and central-memory-like phenotypes described in CTCL, our results are virtually unique among similarly nominated gene signatures [27, 52–54].

### CTCL induces exhibit distinct remodeling of the skin microenvironment

Turning to the broader microenvironments, we next subset and reprocessed skin samples from CTCL, benign inflammatory dermatoses (BID), and healthy control (HC) patients, identifying niche immune cell subpopulations within this compartment (Fig 5a). Initial compositional analyses suggested enrichment of several immune cell populations including B cells and plasmacytoid dendritic cells (pDCs) in CTCL lesions as compared to the other disease states, consistent with previous work [26, 29] (Fig 5b,c). Formal differential abundance testing of these cell types was performed using the *speckle* package [55], but due to the cohort imbalance across these disease states, CTCL and BID samples could not be directly compared. While B cells, plasma cells, and pDCs were among the cell types enriched in CTCL lesions as compared to HC and BID lesions (Fig 5d), these results must be interpreted with caution given the complex library prep and limited cross-cohort design across these defined disease states [29].

**Figure 5:**
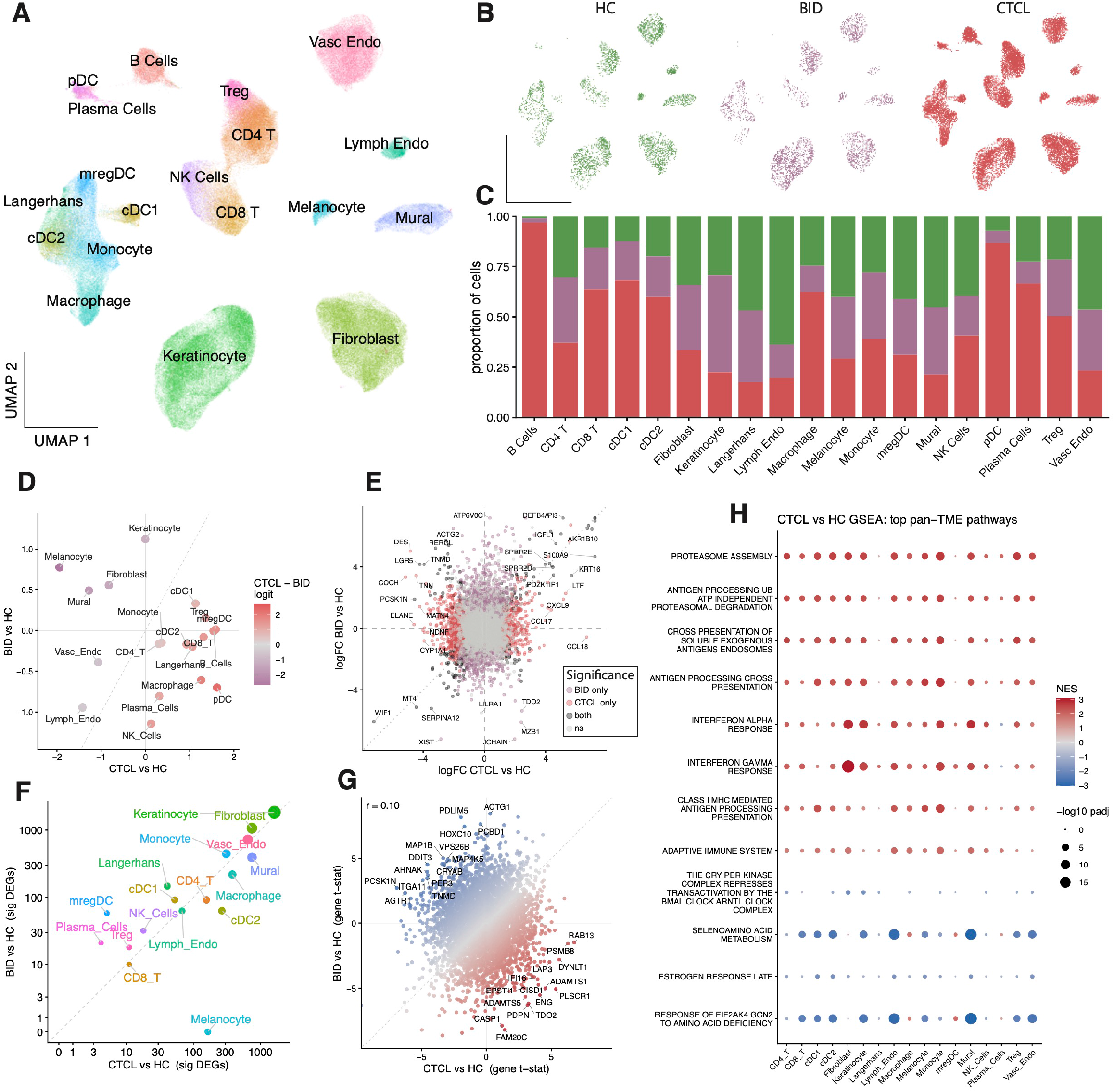
Characterization of microenvironmental changes in CTCL. A. UMAP visualization of the reprocessed skin samples, colored by annotated cell type. B. UMAP visualization of the skin samples colored and split by diagnosis. C. Stacked bar chart of the relative distribution of each identified cell type across each diagnosis following random down sampling. D. Scatter plot depicting the resultant effect sizes of each cell type following differential abundance testing, colored by a scaled difference in these logit-transformed proportions. E. Scatter plot depicting the results of the whole-sample bulk aggregation and DEG testing across diseases of interest, colored according to the significance in each test. F. Scatter plot depicting the number of significant DEGs detected in the cell type-aware pseudobulk testing, colored by annotated cell type. G. Scatter plot comparing the results of the fibroblast pseudobulk testing across disease contrasts of interest, colored by a scaled difference between tested conditions. H. Dot plot of the top recurrent pathways as determined by GSEA of the cell type-aware pseudobulk DEG testing results comparing CTCL vs HC. CD4 T, CD4+ T cells; CD8 T, CD8+ T cells; cDC1, Conventional type 1 dendritic cells; cDC2, Conventional type 2 dendritic cells; Lymph Endo, Lymphatic Endothelial; mregDC, mature regulatory dendritic cells; NK Cells, Natural killer cells; pDC, plasmacytoid dendritic cells; Treg, T regulatory Cells; Vasc Endo, Vascular Endothelial; HC, Healthy Control; BID, Benign Inflammatory Dermatoses; CTCL, Cutaneous T Cell Lymphoma; NS, not significant; DEGs, Differentially expressed genes; GSEA, Gene set enrichment analysis; NES, Normalized enrichment score

Given these limitations, we next assessed these microenvironmental populations for differences in gene expression across disease states. Initially, bulk aggregation of expression across all non-malignant TME cells within each sample was completed to capture combined effects of cellular composition and transcriptional changes associated with disease state. While this pan-TME mixed modeling and GSEA identified significant increases in genes related to immune infiltration, adaptive immune responses, and antigen presentation (Fig 5e, supplementary), this testing cannot identify cell type-specific changes in gene expression.

We next performed pseudobulk aggregation and linear mixed-effects modeling within the identified cell types. This revealed several significant genes across both comparisons of interest, most notably within the stromal cell compartment (Fig 5f). Given the number of significant DEGs within the fibroblast population across both comparisons and the increasing role these cells are believed to play in CTCL [26, 56–58] and cancers, broadly, [59–61] we focused specifically on these results. Interestingly, we found that the underlying gene expression changes associated with CTCL and BID were highly disparate, supporting these as shared inflammatory states with unique underlying transcriptional profiles. Notably, CTCL fibroblasts were uniquely enriched in *RAB13*, *PSMB8*, *CD74*, *PLSCR1*, and *ENG,* among many others (Fig. 5g). Finally, to identify top recurrent pathways across cell types, GSEA was performed on CTCL and BID pseudobulk results. CTCL results revealed significant and coordinated enrichment of many pathways associated with antigen processing and presentation, interferon signaling, and adaptive immune responses, consistent with a chronically inflamed, antigen-presentation-rich microenvironment in CTCL lesions [19, 20, 62, 63] (Fig 5h).

### CTCL lesion types and molecular subtypes drive distinct TME changes

We next examined the role of lesion type and the identified molecular subtypes on the CTCL microenvironments. Restricting the analysis to lesional CTCL samples, we retained all original cell types identified in the original skin analysis (Fig 6a). Similar descriptive compositional analysis suggested the relative specificity of B and plasma cell populations for tumor lesions and melanocytic and keratinocytes for erythrodermic lesions (Fig 6b, c). Formal compositional testing supported these trends, although only the increase in B cells within tumor lesions was determined to be significant (Fig 6d). Interestingly, while the presence of B cells within CTCL lesions has been nominated as a specific finding as compared to HC and BID lesions, we believe that B cell infiltration may be enriched in tumor stage lesions specifically [26, 58].

**Figure 6:**
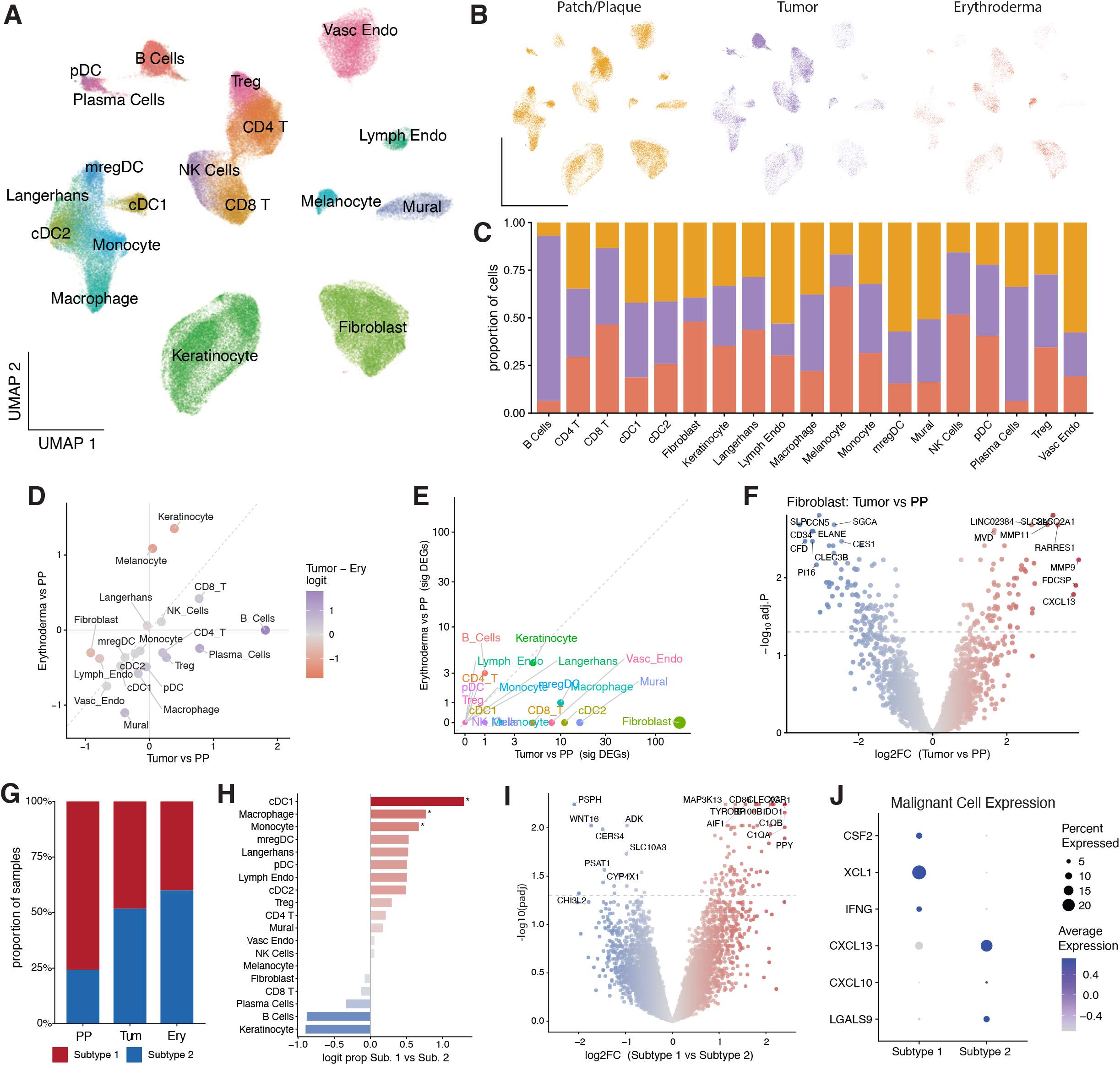
Lesion type and molecular subtype specific TME alterations. A. UMAP visualization of the reprocessed lesional CTCL skin samples, colored by annotated cell type. B. UMAP visualization of the lesional CTCL samples colored and split by lesion type. C. Stacked bar chart of the relative distribution of each identified cell type across each CTCL lesion type following random down sampling. D. Scatter plot depicting the resultant effect sizes of each cell type following differential abundance testing, colored by a scaled difference in these logit-transformed proportions between lesion types. E. Scatter plot depicting the number of significant DEGs detected in the cell type-aware pseudobulk testing between lesion types of interest, colored by annotated cell type. F. Volcano plot of the results of the DEG testing comparing tumor vs patch/plaque fibroblasts. G. Stacked bar chart showing the proportion of samples designated each molecular subtype across each lesion type. H. Bar chart depicting the effect size of each cell type following differential abundance testing. I. Volcano plot of the TME bulk aggregated and DEG testing of the annotated subtypes. J. Dot plots of select top gene expression within the aggregated malignant cell populations for each subtype. CD4 T, CD4+ T cells; CD8 T, CD8+ T cells; cDC1, Conventional type 1 dendritic cells; cDC2, Conventional type 2 dendritic cells; Lymph Endo, Lymphatic Endothelial; mregDC, mature regulatory dendritic cells; NK Cells, Natural killer cells; pDC, plasmacytoid dendritic cells; Treg, T regulatory Cells; Vasc Endo, Vascular Endothelial; DEGs, Differentially expressed genes; PP, Patch/plaque; Tum, Tumor; Ery, Erythroderma

Pseudobulk linear mixed-effects modeling was performed across these distinct lesion types. While we did identify notably fewer significant DEGs in this testing, fibroblasts were once again among the top perturbed cell types (Fig 6e). In total, 179 significant DEGs were identified in the tumor-derived fibroblasts as compared to the patch/plaque-derived fibroblasts (Fig 6f). Among the top changes included significant decreases in expression of *CD34, PI16*, and *CCN5*. Conversely, *SLC2A5*, *MMP11*, *FDCSP,* and *CXCL13* were among the most significantly increased (Fig 6f). We believe that these results represent a shift in the behavior of these cells from a universal fibroblast phenotype towards that of a cancer associated fibroblast (CAF) and an ectopic/tertiary lymphoid structure (TLS) supporting state [60, 64]. Notably, while even fewer significant DEGs were identified across other cell types, both *LAG3* and *HAVCR2* were found to be significantly increased in CD8+ T cells within the tumor lesions.

Finally, we asked whether the identified molecular subtypes of CTCL drove any detectable changes in their TMEs. While both subtypes were present across all defined lesion types, Subtype 2 was relatively more frequent among advanced-stage lesions (Fig 6g). Surprisingly, differential composition testing of these subtypes revealed stark increases in conventional type 1 dendritic cells (cDC1), macrophages, and monocytes in Subtype 1 samples, with relative increases in keratinocytes, B cells, and plasma cells in Subtype 2 samples (Fig 6h). Notably, these results held up even when accounting for lesion type, and pan-TME aggregation and differential expression testing largely supported these as well (Fig 6i). To identify the drivers of these compositional differences, we assessed the expression of a number of canonical ligands and cytokines supporting these disparate immune cell populations. Notably, we found robust expression of these within the original CTCL cell populations, identifying significantly increased levels of *CSF2*, *XCL1*, and *IFNG* in the Subtype 1 cells. Subtype 2 samples had increased expression of *CXCL13*, *CXCL10*, and *LGALS9*, although *CXCL13* was not significant (Fig 6j). While limited, these data suggest that the compositional differences observed between these identified molecular subtypes of CTCL may be coordinated by the intrinsic expression profiles of the CTCL cells themselves.

### Subtype signatures correlate with clinical axes of progression and predict clinical outcomes in an independent cohort

Last, we assessed our results for clinical relevance. Pseudobulk aggregated CTCL profiles were generated for a panel of pertinent CTCL and normal T cell markers (Fig 7a). Pairwise correlation analysis of tissue- and cohort-adjusted residuals was performed across CD4+ phenotypic samples, revealing a complex interplay of genes across disparate functional groups (Fig 7b). To model these changes in gene expression across a clinically-informed and prognostically significant axis of disease progression [65], we performed linear mixed-effects modeling of malignant cell expression across an ordinal encoding consisting of early-stage skin (1), advanced-stage skin (2), and leukemic disease (3). This model identified 3,387 genes with a significant linear trend across this ordered trajectory. Top changes along this encoding included decreases in *IL13*, *CXCR6*, *GZMB*, and *NR4A1* and increases in *AIRE*, *KLF2*, *RASGRP2*, and *TCF7*, consistent with known changes in memory phenotypes (Fig 7c, d) [52–54]. Interested in whether our identified subtypes could be used similarly, linear mixed-effects modeling was performed and revealed that the subtype encoding using Subtype 1 as the starting node resulted in more significant DEGs (n = 3,662) than the reverse as well as the original clinical stage-based ordering. While these results were highly correlated, this suggested that the identified subtypes may capture a known axis of disease progression, despite their independence from clinical stage (Fig 7e).

**Figure 7:**
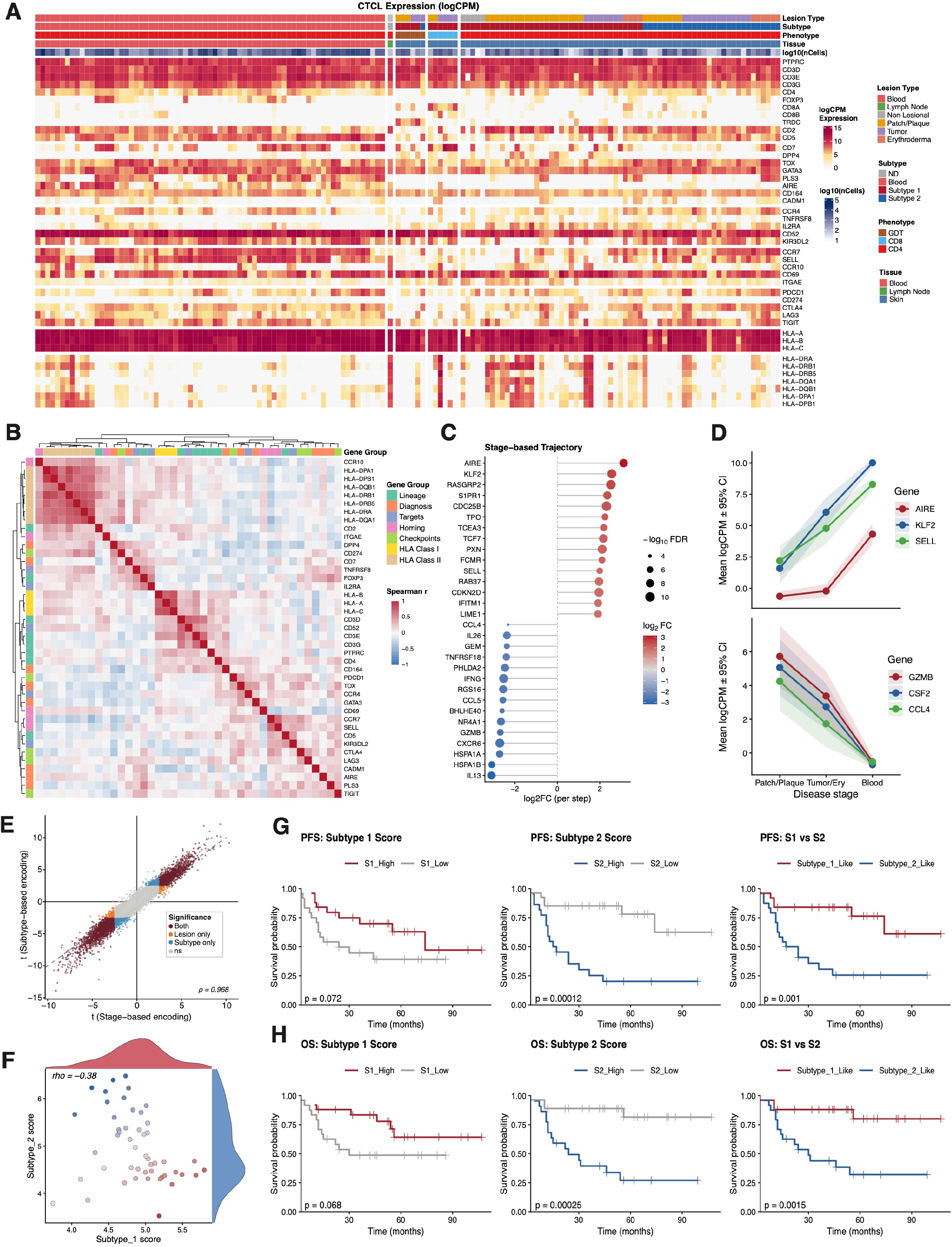
Clinical association and prognostic implications of the molecularly defined CTCL subtypes. A. A heatmap of the log2FC expression values of the aggregated CTCL cells expression across each sample, with sample-level annotations included. B. A heatmap of pairwise, Spearman rank correlation coefficients of the tissue and cohort adjusted gene residuals. C. Lollipop plot of the top increasing and decreasing genes across the clinical stage-based ordinal DEG testing. D. Ribbon plots of the logCPM expression values of select top genes across the clinical stage encoded variables of interest. E. Scatter plot of the results of the clinical stage-based and subtype-based ordinal DEG testing. F. Scatterplot of the subtype-derived gene signature scores, distribution, and Spearman’s rank correlation coefficient. G. Kaplan Meier (KM) plots of the PFS of the independent cohort of patients, split by scoring method and colored according to group classification. H. Kaplan Meier (KM) plots of the OS of the independent cohort of patients, split by scoring method and colored according to group classification. ND, Not determined; GDT, gamma-delta T cell; CD8, CD8+ T cell; CD4, CD4+ T cell; HLA, Human leukocyte antigens; PP, Patch/plaque; Tum, Tumor; Ery, Erythroderma; NS, Not significant; LCT, large cell transformation; NLCT, non-large cell transformation; PFS, Progression-free survival; OS, Overall survival; S1, Subtype 1; S2, Subtype 2

To evaluate the prognostic significance of these subtypes, we downloaded an independent bulk RNA sequencing dataset of advanced-stage mycosis fungoides (MF) patients [17]. Samples were scored according to a gene signature list of the top 25 genes in each subtype. The distribution of these scores was assessed and found to be anti-correlated (rho = -0.38), validating these distinct populations (Fig 7f). To assess the impact of our derived signature scores on known patient outcomes, samples were classified into high/low according to each signature score as well as a composite score representing the difference between the two, and the association of these with progression-free and overall survival (PFS, OS) was assessed using log-rank test (Fig 7g, h). Across all tests, samples scoring high for the identified Subtype 2 gene signature were found to have worse outcomes, with notable significance using either the Subtype 2 or composite scores. We also assessed the prognostic significance of these subtype signature scores using a continuous Cox model unadjusted and adjusted for large-cell transformation status (LCT), a known poor prognostic risk factor. While the results of the continuous Cox model recapitulated the log-rank results, significance was lost when adjusting for LCT status (Supplementary). Because almost no cohorts included in our original analysis note the LCT status of their samples, we were unable to directly assess the association between our assigned subtypes and this clinical feature.

## Discussion

Cutaneous T cell lymphoma is a highly heterogeneous disease process from clinical, histopathologic, immunophenotypic, and molecular perspectives, posing diagnostic and management challenges. Furthermore, due to the complex genetic landscape of CTCL, our understanding of these diseases remains limited, resulting in numerous unmet clinical needs. These include delays in diagnosis, lack of molecularly-informed and targeted treatment options, and risk stratification tools that rely predominantly on limited clinical data (disease extent, laboratory values, age, sex) [66]. By focusing on the downstream changes in expression resulting from these heterogeneous and complex genetic alterations, we hoped to improve our basic understanding of CTCL while identifying novel targets to address these unmet clinical needs.

Toward this end, we generated the largest and most diverse atlas of CTCL scRNA-seq data to date. We combined this with statistically rigorous approaches of analysis, illustrating the ability to elucidate recurrent findings across seemingly diverse cohorts of patients. While many challenges remain in the rigorous and efficient use of publicly available, high-throughput data, cross-cohort integration provides an exciting opportunity to revisit unanswered questions, particularly in the context of rare and heterogeneous diseases such as CTCL.

The single-cell data presented here were originally generated to answer a wide range of open questions in the CTCL field. These efforts have significantly expanded our understanding of CTCL cellular behavior and intra-patient and intra-sample heterogeneity, which is substantial. While unable to fully explore this in the current work, the degree of variation in expression across patients highlights the breadth of transcriptional states present in CTCL. However, applying these data to improve understanding of disease in relation to sample-level metadata, recurrent programs across patients, and clinical outcomes has been challenged by sample size and a failure to properly address pseudoreplication and unwanted sources of variation. While many studies from which this data originated attempted to link cell-specific and CTCL-specific expression to sample-level metadata, here we show that these results largely reflect of the specific patient samples and fail to generalize across a wider cohort.

We sought to address many of these same questions – starting with the basic question of what makes CTCL cells different from their benign counterparts. While the loss of expression of *CD7* and *DPP4* (CD26) and the expression of *TOX*, *PLS3*, and *KIR3DL2* are the most common markers of malignancy and used to help identify these cells for clinical diagnosis, even these are not present in all cases. Using our approach of pseudobulk aggregation and linear mixed-effects modeling, we were able to nominate several thousand significant DEGs, including genes with stronger differential expression than several of these canonical markers. Among the top increased genes included *SUN1* and *MFSD10*, outranking even *TOX*, as well as *HDAC9*, *CD70*, *VPS25*, *KLHL42*, *PSMB3*, *PTHLH*, and *PGM2L1*. Notably, histone deacetylase (HDAC) inhibitors are widely used in CTCL, although many of these are pan-class or multi-class targeting. Consistent with a previous report using assay of transposase accessible chromatin with sequencing (ATAC-seq), *HDAC9* appears to be relatively specifically enriched in CTCL and could represent an opportunity for specific targeting [67]. Conversely, these efforts identified numerous downregulated genes that were more significantly downregulated than even *CD7*. Among these include *GZMK*, *APBB1*, *LINC01550*, *ITGA4*, *TGFBR2*, *PTGER2*, *SATB1*, and *STAT4*. While some of these may represent finer phenotypic differences yet present in the dataset, *TGFBR2* and *PTGER2* may represent similar losses of immunomodulatory signaling by transforming growth factor-beta (TGF-β) and prostaglandin E2 (PGE_2_). Loss of responsiveness to TGF-β is a documented mechanism of resistance and therapeutic vulnerability in CTCL, despite evidence that PGE_2_ may be a growth factor in MF [46, 68]. Finally, *SATB1* appears a particularly intriguing pathogenic candidate gene given its known role in regulating genome architecture, TCR signaling, and hematopoietic stem cell self-renewal, as well as the role of TCR signaling, maintenance of self-renewal, and widespread alterations in genomic structure and stability in CTCL [69–71]. Importantly, while many studies have attempted similar, including using pseudobulk method, these studies have failed to remove disparate phenotypes from their comparator population, making it difficult to assess what changes are a result of differences in normal T cell phenotypes versus related to the underlying malignancy process. While we were unable to identify specific CD4+ T helper (Th) subsets in our data due to the poor sequencing depth and low expression of genes used to discern these populations, we were able to identify this non-Treg CD4+ population for comparison.

We also applied a robust approach to identify and characterize molecularly defined subtypes of CTCL within the skin compartment. Although these subtypes appear to represent phenotypes similar to those classically and more recently described in the CTCL literature, this is the first work to take a statistically robust approach to characterize these subtypes, identifying several hundred novel, significant DEGs between them [27, 52]. As evidenced by this, we show that our derived gene signatures had significant prognostic implications, outperforming previous efforts [27]. Importantly, we did find that these subtype signatures differed significantly by LCT status. When adjusting for this in our assessment of PFS and OS of these patients, our results were no longer significant. Unfortunately, the LCT status of most original single-cell samples was not documented, so the effect that this process has on CTCL cell expression could not be assessed. Given the prognostic significance of LCT, this represents a notable gap in the field and significant opportunity to apply these single-cell technologies for further prognostic refinement [72–74]. Furthermore, LCT status is a binary and subjective interpretation [75, 76]. Despite the association of our subtype signature scores with this, our expression-based scoring system represents an objective and continuous measure that could better reflect continuous risk scores and should be assessed in future work.

Finally, to identify recurrent microenvironmental changes unique to CTCL and across these disparate lesion types and identified molecular subtypes of disease, we applied the same robust modeling to the abundance and expression of normal cell populations within these lesions. While this analysis was complicated by the effects of cohort imbalance, small sample sizes, and disparate sorting prior to library preparation, we identified several significant alterations in the abundance and expression of these TME population across all three contrasts of interest. First, this testing suggested shifts in the composition of the abundance of the identified cell types in CTCL as compared to HC and BID disease states. Additionally, differential gene expression testing revealed stark and CTCL-specific expression changes across numerous cell types in CTCL lesions. Most notably, CTCL fibroblasts were found to have significant increases in *RAB13, PSMB8, PSMB9, PLSCR1, ENG, LAP3,* and *TDO2*, among many others. Many of these genes are implicated in the cancer-associated fibroblast (CAF) phenotype as well as antigen presentation and immunosuppression [59, 64]. GSEA confirmed the enrichment of these pathways in the fibroblasts as well as many of the other cell types, suggesting a highly coordinated, antigen-rich environment in CTCL.

Second, distinct lesion types of CTCL were assessed for differences in cellular abundance and expression profiles. While this testing was limited for the erythrodermic lesions, comparison of the patch/plaque-type lesion to the tumor-type lesions was well powered. Building on previous work identifying B cell infiltration as a specific finding in CTCL lesions, we found a significant association between B cells and tumor lesions. In addition, the identification of *CXCL13* and *FDCSP*, critical genes in TLS formation, expressed by tumor lesion fibroblasts supports the potential development of TLS-like aggregated in these advanced-stage lesions. *CXCL13* likely is responsible for the noted increased presence of B cells in these tumor lesions, driven by binding of *CXCR5*. Originally discovered to be expressed in follicular dendritic cells and intimately tied to peripheral lymphoid organ development, *FDSCP* has since been shown to be expressed in a number of tumors, and here we show the expression of this gene specifically in tumor-derived fibroblasts in CTCL [58, 77–80]. While the number of significant genes was relatively limited across other cell types, CD8+ T cells were found to have significant increases in both *LAG3* and *HAVCR2*, suggesting a potential role of CD8+ T cell dysfunction with these advanced-stage lesions [81–84]. Subsequent investigation revealed increases in a number of additional immunosuppressive genes in these cells including *PDCD1* (PD-1) and *CTLA-4*, although these were not significant [85].

Third, investigation of the identified subtypes revealed significant shifts in the abundance of specific immune cell subsets. Subtype 1 samples were found to be enriched in cDC1, macrophage, and monocyte populations, potentially contributing to the highly inflammatory environment typically found in early-stage lesions, while Subtype 2 samples were found to be enriched in plasma and B cells, although these were not significant [86, 87]. While limited, the lack of identified significant DEGs across TME cell types and identification of large differences among CTCL expression of *CSF2*, *XCL1*, *IFNG*, *CXCL13*, *CXCL10*, and *LGALS9* may support a direct role for CTCL cells in coordinating these changes. *XCL1* is known to specifically recruit cDC1 cells through binding of *XCR1* [88, 89]. Additionally, *CSF2* and *IFNG* are both known to play critical roles in the trafficking and skewing of inflammatory monocytic, macrophage, and granulocyte responses [90, 91]. Conversely, *CXCL13* plays a central role in recruiting B cells and other lymphocytes through interaction with its canonical receptor, *CXCR5* [92]. Finally, while less directly involved in the trafficking of B and plasma cells, *CXCL10* is also a key ligand involved in recruitment of diverse immune cells through binding of *CXCR3,* and *LGALS9* has been implicated in T cell trafficking as well as inhibiting immune responses through binding of TIM-3 and CD44 [93–96].

In summary, our work nominates recurrent expression changes across a diverse cohort of CTCL patients, identifies molecularly defined subtypes of disease within the skin compartment with significant prognostic implications, and illustrates shifts in cell type abundances and expression profiles as they relate to CTCL, distinct lesion types, and the identified molecular subtypes. We hope that these findings will inform future studies in CTCL and to help address the many unmet clinical challenges for these patients.

## Methods

### Study design and data acquisition

Relevant single-cell RNA sequencing (scRNA-seq) samples were identified through a literature search, and the associated public datasets were obtained via query of the Gene Expression Omnibus (GEO), ArrayExpress and Sequence Read Archive (SRA) databases. In total, 244 unique samples spanning 132 patients were obtained for analysis. Samples spanned three tissue compartments – skin (n = 157), peripheral blood (n = 86), and lymph node (n = 1). A Seurat Object was generated from the available filtered feature-barcode matrices for each sample using the *Seurat* package v5.4.0 [97] and the R programming language v4.5.3. The *BPCells* package v0.3.1 [98] was used to store and access expression matrices on disk, while allowing for memory-efficient retrieval as needed.

### scRNA-seq quality control and filtering

Gene-count matrices were generated for each sample, and gene names were updated to approved gene symbols using the HUGO Gene Nomenclature Committee (HGNC) database using the *HGNChelper* v0.8.15 package [99]. The *decontX* v1.4.1 package [100] was used to remove ambient RNA per sample using default parameters. Doublets were identified using the *scDblFinder* v1.20.0 package [101] and predicted doublets were removed. Finally, relevant metadata was unified across cohorts and samples combined into a single Seurat Object. This object was normalized using log-normalization, cell cycling scoring was completed using *Seurat*, and mitochondrial (mt), ribosomal (rb), and hemoglobin (hb) gene percentages were calculated for each cell. Cells with greater than 20% mt, 60% rb, or 5% hb were removed from downstream analysis.

### scRNA-seq preprocessing

From the resultant filtered object, variable features were identified and TCR, BCR, and hemoglobin genes were removed. From this, 100,000 representative cells were sampled using the method “LeverageScore” in the *Seurat* package. Using this subset of cells, variable features were recalculated (TCR, BCR, hemoglobin, and cell cycle genes were removed), the data layer was scaled (vars.to.regress = “cc.total”, “percent.mt”, “percent.rb”), principal component analysis (PCA) was completed, the samples were integrated using the *harmony* v1.2.3 package [102] per sample, and a UMAP was generated using the first 30 harmony-corrected dimensions, n.neighbors =30, and the umap-learn method. Nearest neighbors were calculated using these same dimensions and clusters were identified across a range of resolutions. Major cell type annotations were assigned to these clusters, and the full dataset was projected to the sketch-based integrated results and assigned cell types. Cells assigned a mapping score less than 0.8 were removed and the final object containing 1.57 million cells was saved for downstream analysis.

### T cell receptor clonality processing and clonality definition

Paired T cell receptor sequencing (TCR/VDJ-seq) data, where available, was integrated with the gene expression data using the *scRepertoire* v2.2.1 package [103]. TCR gene usage and cdr3 sequences were assigned on a per cell basis. For cells with multiple TRA or TRB chains detected, the most frequently observed chain was assigned. Clonality was assigned within each individual sample using TCR alpha/beta gene usage. Degree of clonal expansion was calculated for each cell with assigned TCR data using the following cutoffs: Hyperexpanded (0.1 < x <= 1), Large (0.01 < x <= 0.1), Medium (0.001 < x <= 0.01), Small (1e-4 < x < 0.001), and Rare (0 < x < 1e-4). Cells without a detected TCR alpha or beta sequence were assigned Missing (NA).

### Biological sample aggregation

For those samples originating from the same biological source, samples were merged into a single biological sample (orig.merge) for downstream tasks. These include samples which were sorted into their malignant and non-malignant fractions, samples where the dermis and epidermis were separated prior to sequencing, and other enrichment techniques used to artificially separate cells originating from the same biological specimen.

### Sample cell type and clonality proportion calculations

For sample-based proportion analyses, per sample composition was computed as the fraction of each metadata grouping of interest as a function of the total number of cells within that sample. Results are presented on a per sample basis with the proportion of each grouping represented as the respective coloring within the stacked bar chart.

### Cluster and cell type marker identification

Lineage and cluster-based markers were identified using a simple Wilcoxon rank-sum test. The difference in percent expression/detection for each gene and cluster was calculated as pct.1 – pct.2. Top markers were selected based on a combined rank score of log2FC and percent difference expression.

### Malignant cell identification and assignment

Malignant cells were identified and assigned on a per-sample basis using an integrated approach of both gene expression and TCR-seq data, when available. T and ILC cells were subset from each sample and reprocessed using the same workflow as above, without integration. Using known canonical markers of CTCL and cells with detected TCR sequences, cells were assigned “malignant” or “non-malignant” using cluster-based, reduced dimension-based, and TCR-based assignment. These annotations were saved and combined with the original dataset composed of all samples for downstream use.

### CTCL vs reactive T/ILC cell differential gene expression analysis

Differential gene expression analysis at the cell-level was performed using a Wilcoxon rank-sum test across all identified CTCL and background reactive T/ILC cell populations and presented as a function of log2FC (CTCL vs normal) and percent difference in expression (percent CTCL expressing vs percent normal expressing). Select top and CTCL canonical markers were highlighted for illustrative purposes. Following identification of top global DEGs, samples were subset, and per-sample cell-level DEGs were identified using the same Wilcoxon rank-sum test. The number of significant DEGs (adjusted p-value < 0.05 & log2FC magnitude > 0.5) was calculated for each sample and aggregated across genes to assess recurrence across samples. To investigate the effects of technical factors on the number of DEGs (nDEG) detected per sample, we compared the nDEG to the log10-transformed number of CTCL cells detected per sample as well as CTCL-normal composition imbalance (CTCL fraction – 0.5) individually (Spearman) and jointly (linear model). The results of the Spearman rank correlation of nDEG and number of CTCL cells are included in the plot, and a line of the ordinary-least-squares fit with its 95% confidence band as included in the visualization.

### Log2FC shrinkage and aggregation

Per gene log2FC values for each sample underwent adaptive shrinkage using the *ashr* v2.2.63 package [104]. The estimated standard errors (SE) and z-score were supplied under a zero-centered normal mixture prior. This was done to reduce the magnitude of lowly expressed and other low-confidence gene effects towards zero, reducing their influence on downstream aggregation analyses. These results were subsequently aggregated across samples, filtered to only those genes significant in at least 10% of samples, and ordered according to total summation across samples. Samples were ordered by tissue of origin and unbiasedly clustered within these groups. Mean shrunk log2FC values were calculated across samples for which that gene was found to be significant.

### Normal CD4+ T cell isolation

To reduce the effect of normal phenotypic variation in the benign T cell population used for pseudobulk differential gene expression testing, normal T and ILC cells were subset from the full dataset and cells were randomly downsampled using a cutoff of 5,000 cells per sample. These cells were reprocessed, integrated, and clustered, and only those clusters identified as non-Treg CD4+ T cells were retained. We additionally applied stringent filtering at the cell level, removing any cells with detected expression of *CD8A*, *CD8B*, *TRDC*, *FOXP3*, *NCAM1*, or *KLRD1*.

### Pseudobulk aggregation

Throughout the analysis, pseudobulk aggregation was completed across various cell type x sample combinations of interest, with a minimum threshold of 10 cells per pairing. For each of these, the RNA counts layer was aggregated using a simple summation across all gene counts for all cell type x sample combinations of interest, as depicted in Figure 4a. These aggregated “pseudobulk” profiles were normalized and preprocessed using *edgeR* 4.8.2 [105].

### Pseudobulk principal component analysis

Principal component analysis was used to visualize the results of the pseudobulk aggregated profiles across metadata of interest. Aggregated profiles were batch corrected using *limma* v3.66.0 [106]. Tissue and malignancy status were specified as sources of variation to retain in batch correction for the combined CTCL and benign T cell batch correction, while only tissue was retained for the CTCL-specific batch correction. PCA was performed on this batch-corrected matrix, and results were presented across select metadata groupings of interest using the first two principal components (PCs).

### Pseudobulk gene expression variance partitioning

Estimation of sources of variation for each gene across select metadata variables of interest was completed using the *variancePartition* v1.41.4 package [16], quantifying the fraction of expression variance attributable to each variable as well as remaining variance unexplained by the tested variables (residuals).

### Linear mixed-effects pseudobulk differential gene expression testing

Differential gene expression testing on pseudobulk aggregated expression results was completed throughout the analysis via linear mixed-effects modeling using the *variancePartition* package [16]. All testing was completed on TMM-normalized and filtered matrices with TCR and BCR genes removed. All testing modeled the primary contrast of interest as a fixed effect and patient and cohort as random effects, unless noted otherwise. For the CTCL vs CD4+ T cell testing, samples were restricted to CD4+ phenotypic CTCL samples. Tissue was also modeled as a fixed effect. TME pseudobulk testing was completed modeling cohort as a random effect for the CTCL vs HC testing and only the primary contrast of interest for the BID vs HC testing. Lesion type and subtype testing was completed modeling cohort as a random effect.

### Gene set enrichment analysis

Gene set enrichment analysis (GSEA) was performed using the *msigdbr* v26.1.0 [51] and *fgsea* v1.26.2 packages [107] for database query and enrichment analysis, respectively. Enrichment was performed throughout the analysis using descending-ranked gene lists of resultant t-values from the respective DEG testing. Pathways were filtered to those with minSize > 15 and maxSize < 500. FDR correction was applied within each contrast tested.

### Transcription factor inference analysis

Transcription factor (TF) inference analysis was performed using the *decoupleR* v2.16.0 package [108]. Human regulon priors were accessed from the CollecTRI collection [109] using *decoupleR*. TF activity scores were calculated using a univariate linear model with minimum regulon size of 5 targets. P values were adjusted across tests using the Benjamini-Hochberg method.

### Consensus clustering

The batch-corrected expression matrix from the pseudobulk aggregated CTCL profiles was subset to the top 1000 most variable genes and scaled across genes. Robust consensus clustering was performed using the *ConsensusClusterPlus* v1.74.0 package [110] using a random seed. This was completed using 1000 resamples, 80% item sampling, hierarchical clustering, Pearson distance, and average linkage for k values ranging from 2 to 9. Samples were annotated according to the results of the k=4 results.

### TME data preprocessing

Skin samples were subset from the full dataset and CTCL cells removed from the object. Following this, additional stringent filtering was applied using the following criteria: nFeature_RNA > 500, nCount_RNA > 800, percent.mt < 10, percent.rb < 50, percent.rb > 5, cc.total < 0.25. Additionally, samples were randomly down sampled to a maximum of 5000 cells per sample. The resultant object was reprocessed and subsequently split into immune and non-immune fractions for improved annotation.

Each fraction was reprocessed and reintegrated across samples and cohorts using *harmony* [102]. Nearest neighbor analysis and clustering were performed on the harmony-corrected reduced dimensions across a range of clustering resolutions. Low-quality clusters as determined by poor sequencing quality and those containing nearly all cells specific to a single cohort or sample were removed. Finally, these remaining high-quality cells and annotations were saved and these fractions merged into a single object. This object was reprocessed, integrated, and UMAP generated for visualization purposes.

### TME cell type proportions across disease states

The final merged TME object containing both immune and non-immune fractions was randomly downsampled across disease states of interest. The proportion of each disease state within each cell type was calculated and presented as a filled bar chart, with the color corresponding to the relative percent of each grouping of interest per cell type.

### Differential abundance testing

Formal compositional analysis was performed for each TME figure using the *speckle* v1.10.0 package. For each specimen, per-cell-type proportions were calculated and logit-transformed to account for the compositional nature and variance considerations of the data. These transformed proportions were then modeled, and cohort effects were mitigated with a block design using *limma* [106]. The resultant coefficients were moderated with empirical Bayes (eBayes). For the diagnosis and lesion type comparisons, each contrast was made relative to the designated reference. For the CTCL subtype contrast, Subtype 2 was set as the reference, and results are reported using this same convention. These subtype results were also repeated, modeling lesion type as a random effect and included in the supplementary material.

### TME bulk aggregation and gene expression testing

Samples were aggregated across all identified cells using the final, merged TME object in an attempt to identify top DEGs between conditions of interest while retaining the effects of differential cell type composition. Bulk DEG testing was completed using the same *edgeR* preprocessing and *variancePartition* linear modeling. CTCL vs HC testing was completed modeling disease as a fixed effect and cohort as a random effect across all pertinent cohorts. BID vs HC testing was completed only modeling disease as a fixed effect within the single cohort from which the samples originated.

### CTCL subtype ligand expression

CTCL expression of select ligands was calculated across each identified molecular subtype and the percent of cells within each subtype and scaled average expression was calculated for each. Results are presented as a dot plot, split by subtype.

### CTCL clinical marker expression and correlation

A pseudobulk aggregated expression matrix was generated for the CTCL cells from each sample. This matrix was TMM-normalized and logCPM transformed using a prior count of three. This matrix was subset to select markers of interest. Genes were ordered according to functional groups and samples ordered by tissue, malignant phenotype, molecular subtype, and finally lesion type.

### Pseudobulk gene-gene correlation

The above pseudobulk CTCL matrix was filtered to CD4+ CTCL samples from the blood and skin and renormalized and filtered. A linear mixed model was fit on this, modeling tissue as a fixed effect and cohort as a random effect. Model residuals were extracted for the panel genes and pairwise Spearman correlations between genes were calculated across all genes. Genes were unbiasedly clustered using hierarchical clustering and annotated according to their functional groups for visualization.

### Gene expression ordinal trajectory analysis

Samples were assigned an ordinal variable based on lesion type or subtype. Lesion types were assigned as follows: early-stage skin lesions (non-lesional, patch/plaque = 1), late-stage skin lesions (tumor, erythroderma = 2), and blood (leukemic disease = 3). Subtypes were assigned using both possible encodings of the subtypes from the skin (Subtype 1 = 1, Subtype 2 =2; Subtype 2 = 1, Subtype 1 =2), with blood samples encoded as above (3). The *variancePartition* package [16] was used to model gene expression across all three encodings, modeling each trajectory as a fixed effect and cohort as a random effect. Results were aggregated across tests, and a Spearman rank correlation was calculated for each.

### Independent bulk RNA sequencing data acquisition

To assess the association of our identified molecular subtypes of disease in an independent dataset with known annotated clinical data and long-term clinical outcomes, we downloaded an independent bulk RNA sequencing dataset (GSE168508) and associated metadata using the *GEOquery* v2.74.0 package [111]. Following summation of transcript counts across gene symbols, the resultant counts matrix was normalized using logCPM transformation via *edgeR* [105]. Genes were filtered to remove those with logCPM < 1 in more than 10% of samples. Finally, the associated metadata was cleaned and aligned to this normalized expression matrix.

### Subtype signature scoring and prognostic effect scoring

The top 25 genes by t-value were extracted from the subtype gene testing and intersected with the processed independent expression matrix. The mean expression of each gene was calculated across each sample, and a simple mean cutoff was applied to classify samples into high/low signature scores. Additionally, a composite score was calculated as the difference between Subtype 1 and Subtype 2 scores (higher composite score = more Subtype 1-like). The correlation of the subtype-derived scores was assessed via Spearman’s rank correlation. A mean cutoff was used to classify each sample according to this composite score. Using the *survival* v3.8.6 package [112], association of each assignment with progression-free survival (PFS) and overall survival (OS) were assessed by log-rank test and visualized via Kaplan-Meier curves.

### Subtype signature and large cell transformation

To assess the association of our derived subtype signatures with large cell transformation (LCT), a known poor prognostic risk factor in patients with CTCL, samples were scored according to the above protocol and the distribution of these scores was tested between LCT and non-LCT (NLCT) patient samples. The significance of each of these was assessed using a Wilcoxon rank-sum test.

### Univariable continuous Cox modeling

In addition to the mean-split and Cox proportional-hazards regression testing above, we also tested the subtype and composite scores using a continuous Cox model. These results are included in the supplementary and recapitulated the results of the mean-split testing in univariate analysis. Hazards ratios (HRs) in the supplement are reported per 1 standard deviation (SD) of the scaled scores.

### Multivariable Cox modeling adjusted for LCT status

To assess the prognostic significance of our derived scores independent of LCT status, we fit continuous Cox proportional hazards models adjusting for LCT status. Scores were scaled prior to modeling, and HRs are reported per 1 SD of these scores.

## Data Availability

The newly generated single-cell RNA sequencing (scRNA-seq) data generated for this project are deposited to be made publicly available via the Gene Expression Omnibus (GEO) Accession number (GSEXXXXXX). All additional scRNA-seq samples are available via their existing sources from previously published works. Finally, intermediate results used to generate the main figures are publicly available via the GitHub repository: https://github.com/ryanreis333/CTCL_Atlas.

## Code Availability

All code used to generate the intermediate results and figures from the final processed object presented in this publication is deposited at the same public GitHub repository as the intermediate data: https://github.com/ryanreis333/CTCL_Atlas.

## Acknowledgements

We thank the authors and patients from which the publicly available data originated. Data were obtained from a trial funded and sponsored by the National Cancer Institute (NCI) and supported by NCI collaborators, including Merck Research Laboratories.

## Author Contributions

R.J.R., N.B., and A.J. conceived and designed the study. R.J.R and N.B. curated and assembled the datasets and performed the formal analyses. R.J.R. validated the final analyses and generated the final visualizations and supplementary data. P.M.B. and N.B. contributed data and resources. N.B., E.M., V.L., and A.J. provided subject matter expertise and contributed to the final interpretation of the results. A.J. supervised the study and provided resources and funding. R.J.R. wrote the original manuscript. All authors reviewed and edited the manuscript and contributed to and approved the final version.

